# Inflammation and autophagy dysfunction in metachromatic leukodystrophy: a central role for mTOR?

**DOI:** 10.1101/2023.09.14.557720

**Authors:** Zoe Catchpole, Annabelle Hartanto, Tetsushi Kataura, Pawel Palmowski, Andrew Porter, Emma Foster, Kristina Ulicna, Angela Pyle, Robert Taylor, Kate S. Harris, Viktor Korolchuk, Daniel Erskine

## Abstract

Metachromatic leukodystrophy (MLD) is a lysosomal storage disorder typically resulting from biallelic loss-of-function variants in the *ARSA* gene which encodes the lysosomal enzyme, arylsulphatase A, leading to the accumulation of its substrate, sulphatide, and widespread demyelination. Although gene therapy is available for MLD, it is limited by high cost and a narrow window for intervention, which means the development of therapies for MLD remains a key goal. The aim of the present study was to explore disease mechanisms in MLD with a view to identifying novel targets for therapeutic intervention for patients who cannot avail of gene therapy. *Postmortem* globus pallidus and dentate nucleus tissue was obtained from MLD cases (N=5; age 2-33 years old) and compared to age-, sex and ethnicity matched controls (N=5) and studied using discovery proteomics which demonstrated a marked inflammatory response, activation of the mTOR pathway, oxidative stress and metabolic remodelling in MLD cases. Histological analysis of inflammatory markers, including the terminal fragment of complement pathway activation, C3d, and the secreted glycoprotein YKL-40, a commonly used biomarker for inflammation, demonstrated their enrichment in MLD cases. Given that the mTOR pathway plays a key role in supressing autophagy, we next investigated autophagy and identified the accumulation of autophagosomes in MLD cases, consistent with deficient autophagy. Taken together, these findings suggest inflammation and autophagy dysfunction are key processes involved in MLD and that the mTOR pathway could be a novel therapeutic target for MLD.

## INTRODUCTION

Metachromatic leukodystrophy (MLD) is a rare lysosomal storage disorder typically resulting from bi-allelic loss-of-function variants in the ARSA gene, which encodes the lysosomal enzyme arylsulfatase A (ASA) responsible for desulfating sulfatide into sulfate and cerebroside [1]. Impaired ASA activity leads to the accumulation of sulfatide which, through as yet undefined mechanisms, is associated with striking degeneration of white matter [2]. MLD can onset in either late infancy, the juvenile period or adulthood, with late infantile and juvenile forms typically presenting with loss of motor skills followed by cognitive impairment and early death [1]. Adult-onset MLD is less common than late infantile/juvenile-onset forms, but often presents with mild cognitive impairment, psychiatric features often followed by an insidious loss of motor skills [3, 4].

Although no form of MLD can be treated in established disease, a recent gene therapy approach termed Libmeldy© has been introduced in a number of health systems [5]. Libmeldy©, which is based on transfecting haematopoietic stem cells with the functional enzyme and re-implantation into the patient. This approach has been effective in pre-symptomatic late infantile and juvenile cases, but benefits of this approach are much less clear in individuals with established central nervous system symptoms and adult-onset cases [6]. Screening for MLD is currently uncommon, leading to the sub-optimal situation where many patients who could potentially benefit from Libmeldy© are only identified when a sibling succumbs to treatment-limiting symptoms. In addition, recent evidence suggests haematopoietic stem cell-based therapies ameliorate leukodystrophy but are less effective at treating a more insidious neuronal pathology which may lead to later cognitive symptoms [7]. Therefore, taken together, there are compelling reasons to continue developing new therapies for MLD.

MLD is most commonly caused by pathogenic variants in *ARSA*, though can also result from pathogenic variants in *PSAP*, which encodes the activator protein saposin B [8]. ASA activity as measured in leukocytes is broadly associated with MLD sub-types, as it is lowest in younger-onset cases and higher in later-onset cases, though there are a number of reports of individuals with low levels of ASA activity who remain clinically and pathologically unaffected by MLD [2, 9]. Arsa deficient mice do not exhibit leukodystrophy unless the sulfatide-synthesising enzyme, *Gal3st1*, is also over-expressed, suggesting abundance of sulfatide deposition rather than Arsa deficiency alone leads to leukodystrophy in MLD [10]. Studies of peripheral tissues have demonstrated a strong correlation between sulfatide levels in cerebrospinal fluid or sural nerve and electrophysiological evidence of neuropathy in paediatric MLD patients [11]. Taken together, abundance of sulfatide is associated with key pathological changes associated with MLD; however, the mechanisms through which sulfatide induces key pathological changes in MLD are not known. Therefore, there are pressing reasons to interrogate mechanisms underlying demyelination and neurodegeneration in MLD, with a view to therapeutically attenuating these degenerative processes.

The present study sought to use a combination of post-mortem tissue and cellular models to identify biological pathways associated with MLD that could be therapeutically targeted in individuals who cannot benefit from gene therapy.

## METHODS

### Tissue acquisition and preparation

Ethical approval for this study was granted by Newcastle University Ethical Review Board (Ref: 18241/2021). Frozen and formalin-fixed tissue from the globus pallidus and dentate nucleus was obtained from the University of Maryland Brain and Tissue Bank through the NIH NeuroBioBank from MLD cases (N=5) and age-, sex- and ethnicity-matched controls (N=5; Table 1). MLD diagnosis was confirmed in all cases by enzymatic assay to detect deficient ASA activity (Table 1). The globus pallidus and dentate nucleus were selected due to their more limited involvement in MLD, as incorporating severely degenerated regions would be likely to alter the proteome in favour of alternate cell types rather than showing pathological changes to the normal cellular population.

**Table 1:** Demographic information on cases included in the present study.

| Case ID | Diagnosis | Age at death (years) | Sex | Ethnicity | Genetic diagnosis | Cause of death | Dentate | Globus pallidus |
| --- | --- | --- | --- | --- | --- | --- | --- | --- |
| *MLD* |  |  |  |  |  |  |  |  |
| 6011 | MLD | 2 | F | Black | c.1130_1132delTCT / c.465+1G>A | Complications of disorder | X | X |
| 5505 | MLD | 3 | M | White | c.847G>T / c.1010A>T | Complications of disorder | X | X |
| 5509 | MLD | 6 | M | White | c.465+1G>A (Hom) | Complications of disorder | X | X |
| 5785 | MLD | 17 | F | White | c.1283C>T (Hom) | Complications of disorder | X | X |
| 6442 | MLD | 33 | M | Black | c.465+1G>A / c.1283C>T | Complications of disorder | X | X |
| *Controls* |  |  |  |  |  |  |  |  |
| 1275 | Control | 2 | F | Black |  | Acute myocarditis |  | X |
| 1791 | Control | 2 | F | Black |  | Drowning | X | X |
| 5608 | Control | 3 | M | White |  | Ventricular septal defect |  | X |
| 4446 | Control | 3 | M | White |  | Blunt force injuries of the head and torso | X |  |
| 6032 | Control | 4 | M | White |  | Head and neck injuries | X |  |
| 1500 | Control | 6 | M | White |  | Multiple injuries from a motor vehicle accident |  | X |
| 1279 | Control | 17 | F | White |  | Multiple injuries | X | X |
| 6059 | Control | 33 | M | Black |  | Fentanyl intoxication and atherosclerotic CVD | X | X |

Formalin fixed tissues were processed for paraffin wax embedding and cut into 5μm sections and mounted onto glass slides for histological staining and immunohistochemical staining. Frozen tissue for discovery proteomics analysis was prepared by homogenising approximately 20mg in 5% SDS with an ultrasonic probe. Protein concentration was assessed using Micro BCA™ Protein Assay Kit (Pierce™) and 50ug of total protein used for processing and trypsin digestion following S-trap micro protocol. Volume was adjusted to 20ul with S-trap lysis buffer (5% SDS, 50 mM TEAB pH 8.5), then samples were reduced (50mM DTT, 30min, 60C), alkylated (100mM Iodoacetamide, 30min, RT, dark) and acidified (2.5% phosphoric acid). Subsequently samples were loaded onto S-Trap™ micro spin columns (Protifi) in S-trap binding buffer (100 mM TEAB in 90% methanol) and further processed according to manufacturer’s recommendations (trypsin digestion for 1.5h at 47C). Resulting peptide mixture was dried in a vacuum concentrator, redissolved 2%Acetonitrile, 0.1% Formic acid and stored frozen for further analysis. Samples for immunoblot analysis were homogenised in 0.2M TEAB (Sigma) supplemented with protease inhibitors (cOmplete, Roche) with an ultrasonic probe. Protein concentration of samples was then determined with the BCA assay (Invitrogen).

### Identification of pathogenic variants in ARSA

Primer pairs flanking the variant of interest in ARSA (NM_000487.6) were custom designed using primer3 (http://primer3.ut.ee/). Patient DNA was amplified in a standard PCR reaction using GoTaq (Promega). PCR products were purified using Exonuclease I and FastAP (ThermoFisher Scientific) and then Sanger sequenced (BigDye® Terminator v3.1 Sequencing Kit, ThermoFisher Scientific) on an Applied Biosystems 3500 Genetic Analyzer (ThermoFisher Scientific). Data were analysed using SeqScape v2.7 (Applied Biosystems).

### Histological staining

5μm sections from the dentate nucleus and globus pallidus were dewaxed in Histoclear and rehydrated through a graded series of ethanol solutions until water, with the exception of sections to be stained with the alcohol-based stain luxol fast blue (LFB), which were taken to 70% ethanol. Sections were then histologically stained with LFB followed by haematoxylin and eosin (H&E) to visualise myelin and general structure, respectively. Further sections were also stained with periodic acid Schiff (PAS) to identify PAS-positive accumulations indicating glycoproteins and glycolipids, and counter-stained with haematoxylin. To quantitatively evaluate demyelination, LFB/H&E sections were scanned at 10x magnification on an Axioscan-7 slide scanner (Zeiss).

For immunohistochemistry, heat-mediated antigen retrieval in citrate buffer pH6 was performed to unmask epitopes, prior to incubation in primary antibodies (anti-YKL-40, Abcam ab77528, 1:1,000; anti-C3d, Agilent A0063, 1:1,000). Sections were incubated in primary antibodies for one hour at room temperature prior to washing and probing with the universal probe and polymer-HRP components of the Mach 4 Polymer kits (CellPath, PBC-M4U534L). Antibody binding was visualised with DAB and then counter-stained with haematoxylin. Sections were then imaged on an AxioImager microscope (Zeiss) and images were imported into FIJI/ImageJ (NIH, USA) and percentage area immunoreactive for DAB was quantified.

### Discovery proteomics

Samples were analysed by LC−MS/MS using an Ultimate 3000 Rapid Separation LC (RSLC) nano LC system (Thermo Corporation) coupled with e Thermo Scientific™ Q Exactive™ HF hybrid quadrupole-Orbitrap mass spectrometer. 1ug of peptide mixture was loaded, first onto 300μm x 5mm C18 PepMap C18 trap cartridge (Thermo Fisher Scientific) at a flow rate of 10 μL min−1 maintained at 45 °C and then then separated on 50 cm RP-C18 μPAC™ column (PharmaFluidics), using a 100min gradient from 99 % A (0.1% FA in 3% DMSO) and 1% B (0.1% FA in 80% ACN 3% DMSO), to 35 % B, at a flow rate of 300 nL min−1. The separated peptides were then injected into the mass spectrometer via Thermo Scientific μPAC compatible EasySpray emitter at the Ion Transfer Tube temperature of 300oC, spray voltage 2000V and analysed using data independent (DIA) acquisition. The total LCMS run time was 150min. For the full scan mode, the MS resolution was set to 120000, with a normalized automatic gain control (AGC) target of 1e6, maximum injection time of 60ms and scan range of 300−1650 m/z. DIA MSMS were acquired with 50, 12 windows covering 400-1000 m/z range, at 30000 resolution, AGC target set to 1e6, maximum injection time of 55ms and normalized collision energy level of 27%.

The acquired data has been analysed in Spectronaut version 16 (Biognosis) against human proteome database (Uniprot 3AUP00000564-2022.10.20) combined with common Repository of Adventitious Proteins (cRAP), with default settings (Proteotypicity filter: Only Protein Group Specific, Major group quantity: Sum peptide quantity, Minor group quantity: Sum precursor quantity, FDR<1% for both peptides and proteins). The results were then exported as a text file, reformatted using in house written R scripts and further processed in The Perseus software platform version 1.6.15.0 [12]. Pathway analysis was performed on proteins differentially expressed in the MLD group using the DAVID Bioinformatics Resource [13]. For proteomics pathway analysis, the entire identified proteome was used as proteomic background for pathway analysis.

### Immunoblot analysis

For SDS-PAGE/western blot analysis, approximately 10μg of total protein was loaded onto 5-12% Bis-Tris gels with 1mm pore size and electrophorised with either MOPS or MES running buffer in an Invitrogen NuPAGE MIDI tank at 200V, 3000mA, 350W for 40 minutes. Proteins were then transferred onto nitrocellulose membranes using an iBlot2 semi-dry transfer system (Invitrogen) at 20V for 9 minutes, prior to probing with antibodies of interest.

Membranes were blocked in 10% normal goat serum in TBS-T for one hour at room temperature. Primary antibodies (anti-p62, Abcam ab207305, 1:1,000; anti-LC3B, Abcam ab48394, 1:200) were made up in 10% normal goat serum in TBS-T and incubated overnight at 4°C. After washing, membranes were then incubated in appropriate secondary antibodies (goat anti-rabbit-HRP, Invitrogen G21234, 1:10,000; goat anti-mouse IgG_1_, Invitrogen PA174421, 1:5,000) for 30 minutes at room temperature before washing and enhanced chemiluminescence with (SuperSignal™ West Femto Maximum Sensitivity Substrate, Thermo Fisher). Blots were then imaged on an iBright CL1500 system. Sections were then washed in deionised water for 30 minutes and incubated in Ponceau S solution, which was then imaged on an iBright CL1500 system. Analysis of membranes was performed using FIJI/ImageJ (NIH, USA), with area under curve determined for individual protein bands and total protein per lane or dot determined with greyscale Ponceau S images. Abundance of target proteins was determined by normalising target protein to total protein per lane and compared between control and MLD cases.

## RESULTS

### Histological staining

Qualitatively, H&E staining of the dentate and globus pallidus demonstrated ballooned neurons, and these were variably PAS- or LFB-positive, and more obvious in juvenile/adult -onset than late infantile cases (Figure 1). MLD cases had clear demyelination on LFB staining, and this was more marked in late infantile compared to juvenile/adult -onset cases (Figure 1). Quantitative analysis demonstrated a significant 76.3% reduction in LFB staining in the cerebellar white matter of MLD cases (t=4.469, df=6, p=0.004) but no significant difference in LFB staining between groups in the globus pallidus (p=0.284).

**Figure 1.**
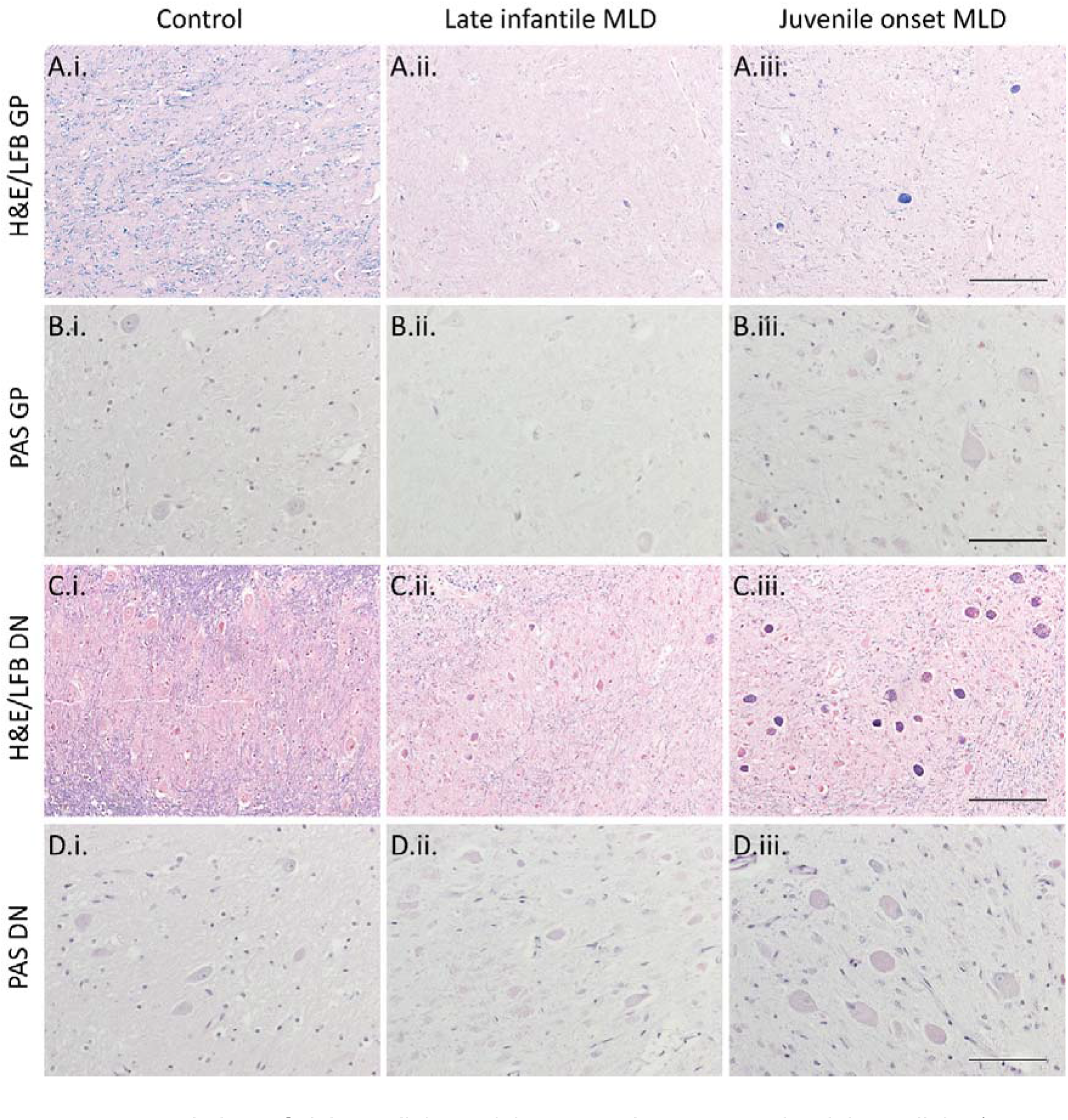
Neuropathology of globus pallidus and dentate nucleus in MLD. The globus pallidus (A.i.-B.iii) and dentate nucleus (C.i.-D.iii.) showed striking demyelination as evidenced by reduction in LFB myelin staining (blue; A.i.-A.iii. & C.i.-C.iii.). Ballooned neurons filled with LFB-positive deposits were observed in MLD cases and especially pronounced in those from juvenile-onset cases (A.iii. & C.iii). Deposits in ballooned neurons were variably PAS-positive (fuchsia; B.iii. & D.iii.), but were especially prominent in dentate nucleus neurons (D.iii.). Scale bars = 200μm (A.i.-A.iii. & C.i.-C.iii.). 100μm (B.i.-B.iii. & D.i.-D.iii.).

### Discovery proteomics identifies inflammation, mTOR activation and metabolic remodelling in MLD

Discovery mass spectrometry identified a total of 5,502 proteins across all samples. To identify differences between control and MLD cases, Perseus software was used with false discovery rate (FDR) correction to attenuate false positives from multiple comparisons [12]. This analysis demonstrated 1,355 proteins were altered in MLD compared to control in the dentate nucleus, of which 533 were down-regulated and 822 were up-regulated. In the globus pallidus, 1,310 proteins significantly altered in MLD compared to control, of which 689 were down-regulated and 621 up-regulated. Pathway analysis using DAVID demonstrated up-regulation of lysosomal components, inflammation and the mTOR pathway in MLD (Table 2). In contrast, down-regulated pathways were less consistent across both regions in MLD compared to controls, though reductions in protein folding and endosomal trafficking were observed in both regions (Table 2).

**Table 2:** Pathway analysis data from dentate nucleus demonstrating pathways altered in MLD dentate nucleus.

| Cluster | GO/KEGG/reactome pathways | Enrichment |
| --- | --- | --- |
| Lysosome | Lysosome, lysosomal lumen | +11.92 |
| Actin | Actin-binding, actin cytoskeleton, actin filament-binding | +3.20 |
| Inflammation | IL1-mediated signalling pathway, TNF-mediated signalling pathway, antigen processing and presentation of exogenous peptide via MHC Class 1, NF-kappaB signalling, proteasome | +3.10 |
| mTOR | Positive regulation of TOR signalling, cellular response to amino acid stimulus, TORC1 signalling, mTOR signalling pathway | +2.21 |
| Pentose-phosphate shunt | Pentose-phosphate pathway, pentose-phosphate pathway (oxidative branch) | +2.02 |
| Endosome | Clathrin-dependent endocytosis, coatamer/clathrin adaptor appendage (lg-like subdomain), endolysosome membrane, clathrin-coated endocytic vesicle | -1.96 |
| Chaperone | Chaperonin-containing T-complex, TCP-1-like chaperonin intermediate domain, chaperonin Cpn60/TCP-1, protein folding | -3.25 |
| TCA cycle | Citrate cycle (TCA), tricarboxylic acid cycle, carbon metabolism, pyruvate metabolism | -3.58 |
| Oxidative phosphorylation | Oxidative phosphorylation, aerobic respiration, mitochondrial ATP synthesis, respiratory chain, mitochondrial respiratory chain complex I assembly | -3.65 |
| Mitochondria | Mitochondrion, mitochondrial matrix | -5.43 |

### Neuroinflammation in MLD

Discovery proteomics indicated up-regulation of a large number of immune-associated proteins (Figure 2A); therefore, we sought to determine whether MLD cases manifested histological evidence of inflammation. We evaluated C3d, a terminal component of the innate immune complement pathway and biomarker of complement-mediated inflammation, in addition to YKL-40, a glycoprotein secreted by macrophages and neutrophils that is considered a marker of chronic inflammation [14, 15]. Histological evaluation of C3d and YKL-40 in the dentate nucleus identified they were primarily expressed in glial cells in late infantile MLD cases, in contrast to neuronal expression in juvenile-onset MLD cases and virtually absent expression in control cases (Figure 2B.i. – C.iv.). C3d and YKL-40 were both significantly higher in abundance in MLD compared to control dentate nucleus (Figure 2B.iv. & C.iv.). Similar patterns were observed in the globus pallidus (Figure 3).

**Figure 2.**
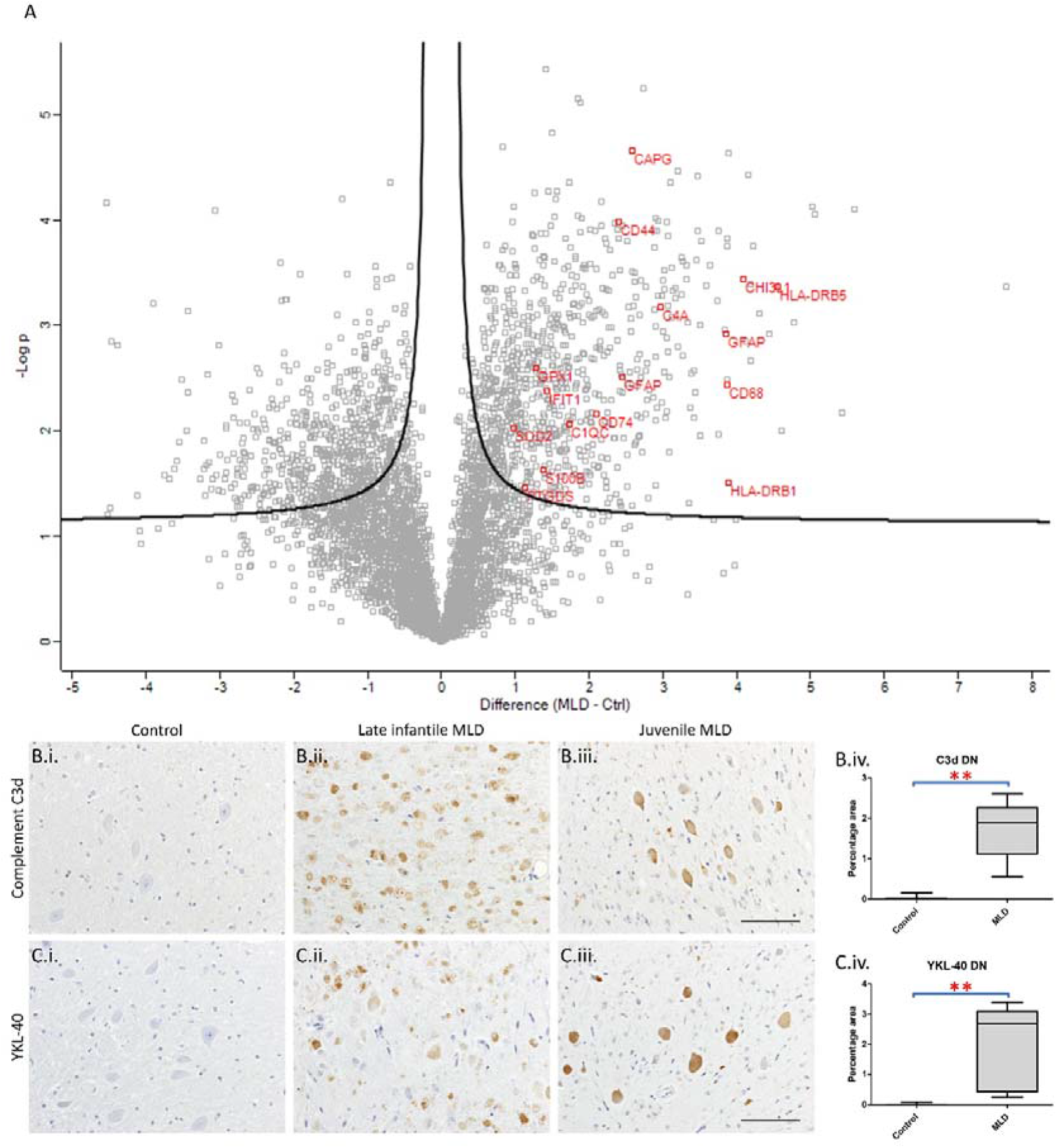
Inflammation in MLD dentate nucleus. Volcano plot demonstrating proteomic differences between MLD and control cases, with select immune/inflammation proteins highlighted in red (A). Histological analysis demonstrated increased Complement C3d (B.i.-B.iv.) and YKL-40 (C.i.-C.iv.) in MLD cases compared to controls. Notably, late infantile MLD cases manifested C3d and YKL-40 immunoreactivity predominantly in swollen glial/macrophage cells (B.ii. & C.ii.) whilst these markers were predominantly neuronal in juvenile-onset cases (B.iii. & C.iii.). **p<0.01. Scale bars = 100μm.

**Figure 3.**
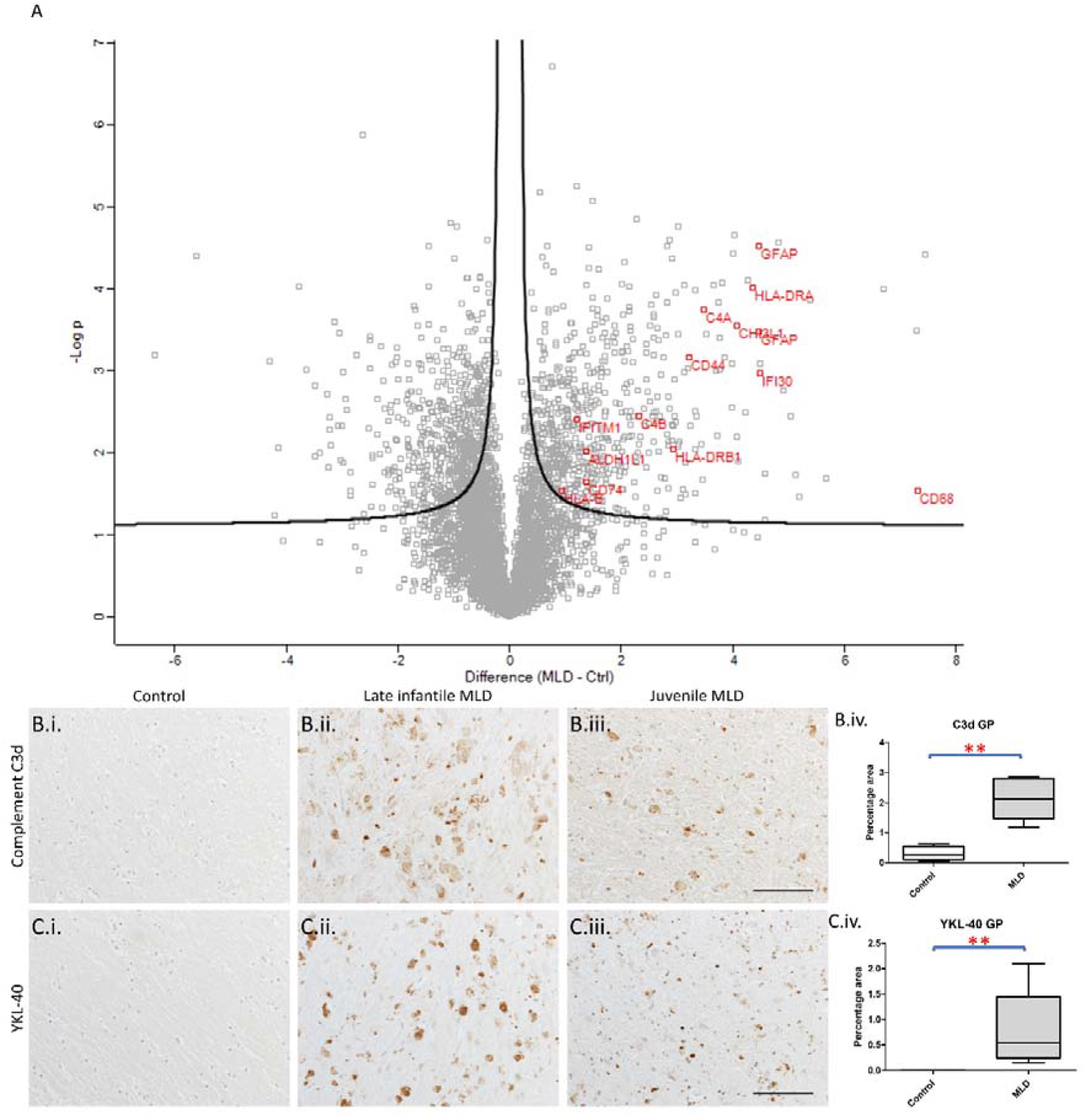
Inflammation in MLD globus pallidus. Volcano plot demonstrating proteomic differences between MLD and control cases, with select immune/inflammation proteins highlighted in red (A). Histological analysis demonstrated increased Complement C3d (B.i.-B.iv.) and YKL-40 (C.i.-C.iv.) in MLD cases compared to controls. Notably, late infantile MLD cases manifested abundant C3d and YKL-40 immunoreactivity, often in swollen glial/macrophage cells (B.ii. & C.ii.), whilst these markers were predominantly in normal appearing microglia and neurons in juvenile-onset MLD (B.iii. & C.iii.). **p<0.01. Scale bars = 100μm.

### Deficient autophagy in MLD

Proteomic analysis identified up-regulation of positive regulators of the mTOR nutrient-sensing pathway, particularly in dentate nucleus. The mTOR pathway plays a critical role in negatively regulating the cellular catabolic process of autophagy, a process which maintains metabolic homeostasis by recycling sub-cellular components [16]. Dysfunction of autophagy has been previously demonstrated to induce metabolic deficits leading to cellular death, and thus the dysfunction of this process has critical and detrimental effects on cellular health [17-19].

Proteomic analysis of the dentate nucleus identified increased abundance of a number of positive regulators of the mTOR pathway in MLD, including a number of components of the Ragulator complex, an essential mechanism for mTOR activation [20]. To determine whether autophagy is impaired in MLD dentate nucleus, western blot analysis of the autophagosome markers p62 and LC3-II was performed on brain lysates, and this identified increased p62 and LC3-II in MLD tissues, consistent with the accumulation of autophagosomes in the context of deficient autophagy (Figure 4).

**Figure 4.**
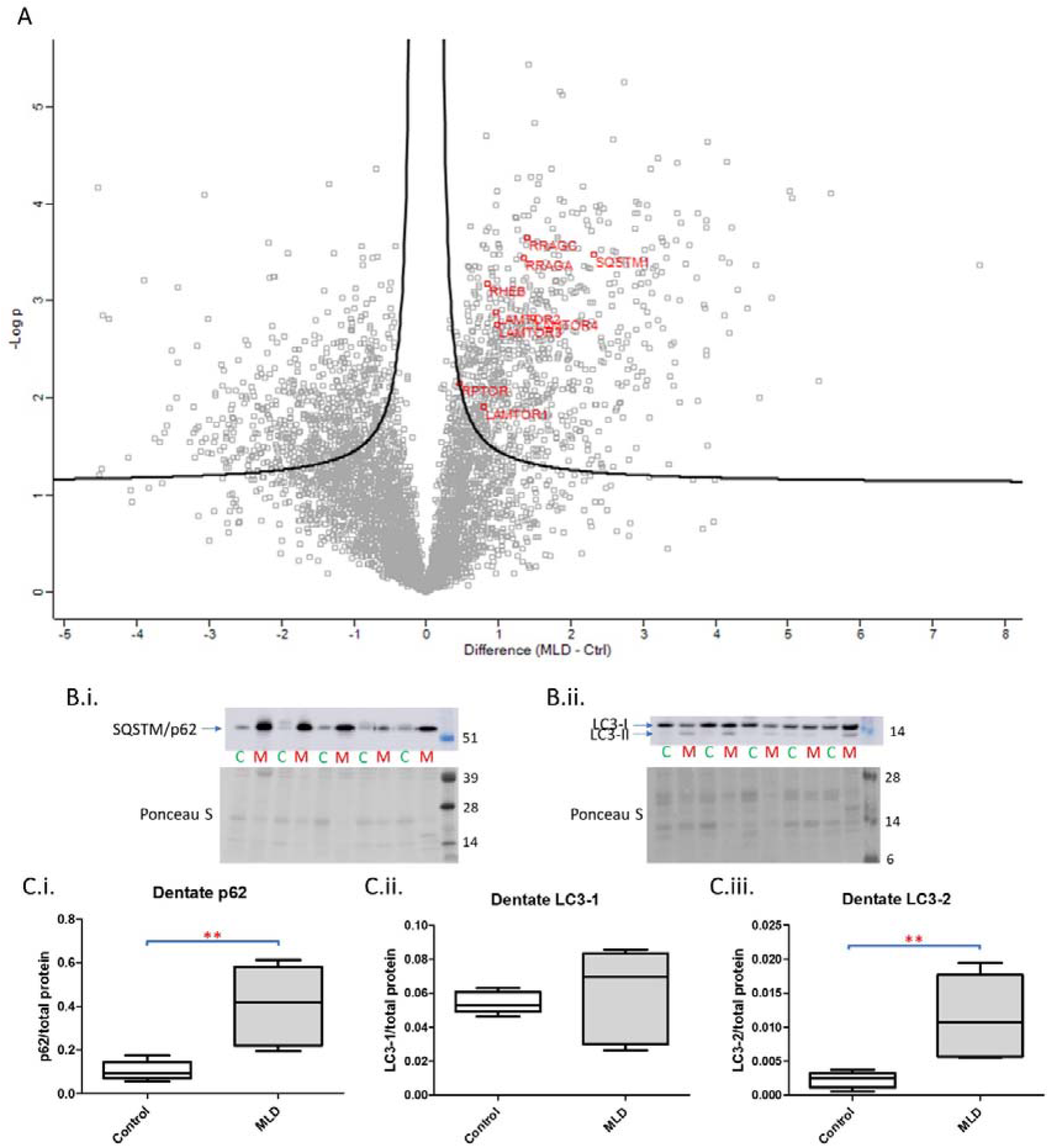
mTOR and autophagy in MLD dentate nucleus. Volcano plot demonstrating proteomic differences between MLD and control cases, with select mTOR activators and autophagosome markers highlighted in red (A). Western blot analysis of the autophagosome markers p62 (B.i.) and LC3-II (B.ii.) demonstrated significant different increases in p62 (C.i.) and LC3-II (C.iii.) in MLD compared to control. **p<0.01.

The globus pallidus showed some evidence of increased autophagosome markers (e.g. SQSTM1) and positive regulators of mTOR (MIOS [21], RRAGA [22] and KLHL22 [23]) in MLD (Figure 5). In addition, decreased abundance of positive regulators of autophagy (e.g. MAP2K1 [24], CHMP5 [25] and HTT [26]) was observed in MLD cases compared to controls (Figure 5). Western blot analysis of p62 and LC3-II once again demonstrated increased abundance of both in MLD, consistent with deficient autophagy and corresponding accumulation of autophagosomes (Figure 5).

**Figure 5.**
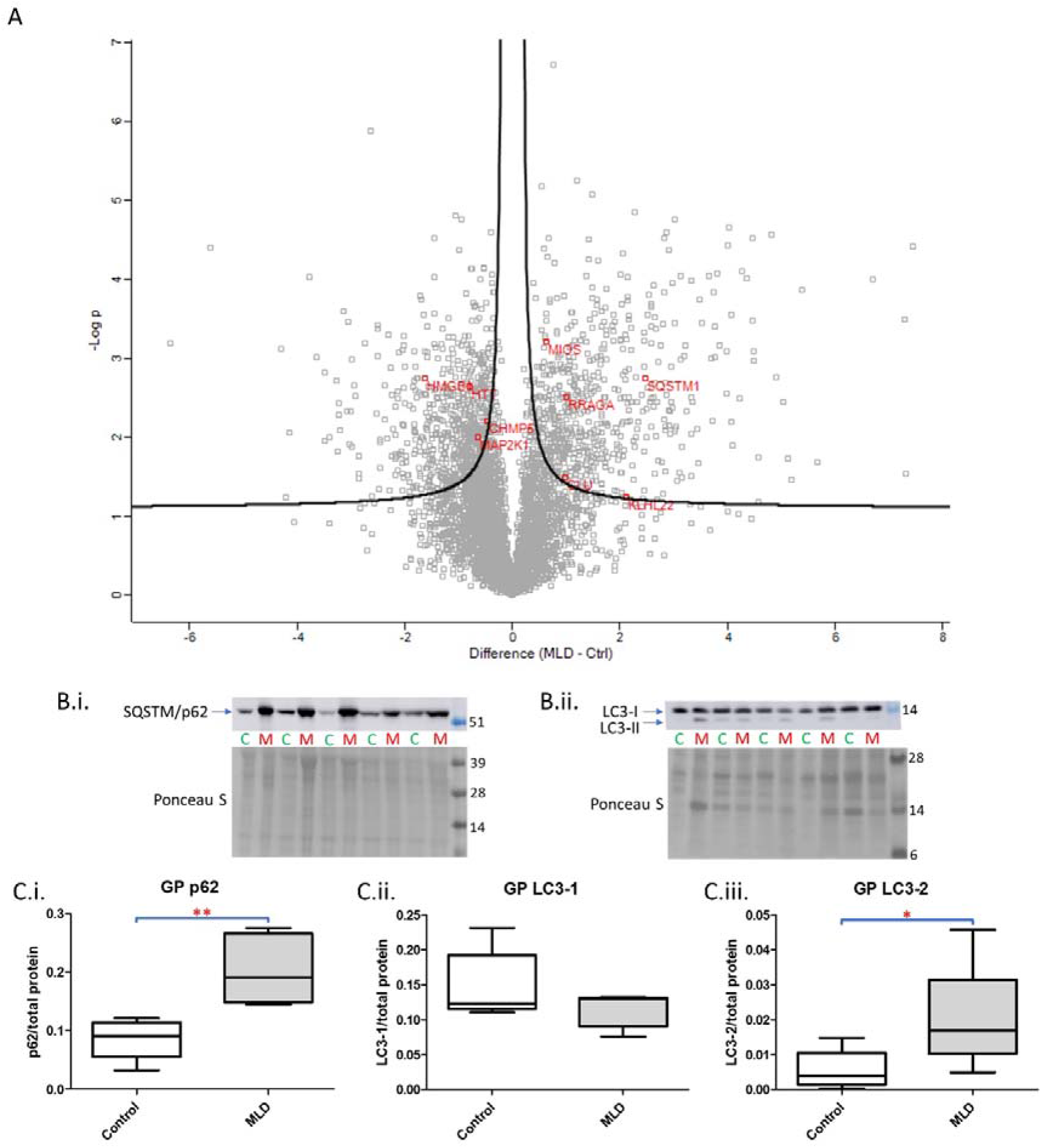
mTOR and autophagy in MLD globus pallidus. Volcano plot demonstrating proteomic differences between MLD and control cases, with select mTOR activators, autophagy activators and autophagosome markers highlighted in red (A). Western blot analysis of the autophagosome markers p62 (B.i.) and LC3-II (B.ii.) demonstrated significant different increases in p62 (C.i.) and LC3-II (C.iii.) in MLD compared to control. *p<0.05, **p<0.01.

## DISCUSSION

In the present study, *post-mortem* MLD brain tissue was investigated using a combined discovery proteomics and neuropathological approach, which identified wide-ranging abnormalities in MLD, including neuroinflammation and deficient autophagy. Proteomics and western blot analysis indicated up-regulation of the mTOR pathway, which is associated with both inflammation and deficient autophagy, which may suggest the mTOR could be a novel therapeutic target to attenuate both autophagy dysfunction and inflammation in MLD.

Previous studies have identified evidence of inflammation in MLD, using cerebrospinal fluid [27] and on neuroimaging [28], leading to speculation that neuroinflammation could play a central role in the pathogenesis of MLD [29]. A previous study in an MLD mouse model demonstrated that neuroinflammation is associated with the degree of sulfatide storage and that attenuation of inflammation with simvastatin reduced demyelination and led to phenotypic improvement [30]. The latter study indicates that inflammation may play a central role in the pathogenesis of MLD. Nevertheless, the key drivers of inflammation in MLD remain unknown, in part as previous *in vitro* studies have not shown a consistent association between sulfatide abundance and inflammation [31, 32]. To the best of our knowledge, the present observations are the first of neuroinflammation as a key aspect of MLD using *post-mortem* brain tissue.

Autophagy dysfunction has previously been reported in a number of lysosomal storage disorders, including Pompe, Gaucher, Danon and Batten disease [33]. We have previously demonstrated deficient autophagy in Niemann-Pick Type C1 [34]. However, to the best of our knowledge, no study has yet demonstrated deficient autophagy in MLD. The presently reported findings of mTOR pathway activation in MLD could underlie both deficient autophagy and inflammation, given mTOR has a key role in regulating inflammation [35]. A recent study in a mouse model of the related condition Krabbe disease demonstrated that inhibition of the mTOR pathway with rapamycin enhanced autophagy, reduced inflammation and attenuated leukodystrophy [36]. These findings are consistent with our observations in Niemann-Pick Type C1, where neurodegeneration was reduced by enhancing autophagy through an effect on NAD levels [18]. The association between neuroinflammation, autophagy dysfunction and mTOR activation in MLD, combined with previous findings demonstrating a beneficial effect on these parameters by activating autophagy, could lead one to speculate that enhancing autophagy and/or inhibiting mTOR activation could be an effective strategy in MLD.

In addition to the central findings of inflammation, autophagy dysfunction and mTOR activation in MLD cases, we also noted qualitative differences between late-infantile and juvenile-onset MLD cases. Notably, we observed a more pronounced glial response in late-infantile MLD cases, comprising the infiltration of morphologically atypical glial cells that were immunoreactive for inflammatory markers. In contrast, juvenile-onset cases had obviously ballooning of neurons, likely indicating accumulation of storage materials, and these were typically reactive for inflammatory markers. Although YKL-40 and C3d are typically associated with glial cells, a previous study has reported C3d in neurons following traumatic head injury, where it appeared to prime neurons for synaptic pruning, and it is thought to label chronically stressed neurons [37, 38]. It is not clear why glial activation is primarily observed in late-infantile MLD, compared to more neuronal pathology in juvenile-onset MLD, but one potential explanation relates to the higher abundance of sulfatides observed in MLD cases under the age of 10 years old compared to those older than 10, presumably related to higher synthesis of sulfatides in early life to support myelination [39]. Previous studies have suggested sulfatide has pro-inflammatory qualities and, therefore, marked gliosis and inflammation in late-infantile cases could be the result of inflammation secondary to sulfatide accumulation [31]. It is not clear why lower sulfatide in later life would underlie a neuronal, rather than glial, pathology in MLD; however, one could speculate that this could relate to a more insidious neuronal pathology that is consistent with marked grey matter atrophy that progresses over time and is more marked in older cases with MLD [40].

Strengths of the present study include the use of MLD brain tissue from individuals with confirmed diagnoses, the use of discovery proteomics to identify disease mechanisms agnostically, and the confirmation of putative pathogenic pathways using histology and/or immunoblotting. Nevertheless, a limitation of the study is the small number of cases, the result of the rarity of MLD and lack of tissue availability. Furthermore, as with all *post-mortem* studies, the cases used were typically end-stage and thus the sequence of pathological changes cannot be inferred, even though we tried to attenuate this confounder as much as possible by selecting regions with minimal degenerative changes.

In conclusion, this study provides neuropathological evidence for a central role for neuroinflammation and autophagy dysfunction in MLD. Given that proteomics data implies increased levels of positive regulators of mTOR activation, and the role of mTOR in sustaining inflammation and reducing autophagy, future studies could investigate whether mTOR inhibition is therapeutically beneficial in MLD cases who cannot benefit from gene therapy.

## ACKNOWLEDGEMENTS

Funding for this study was received by the Alzheimer’s Research UK Northern Network. DE is funded by an Alzheimer’s Research UK Senior Fellowship (ARUK-SRF2022A-006). TK was supported by Fellowships from Uehara Memorial Foundation and the International Medical Research Foundation. VIK was supported by a VitaDAO/Molecule academic partnership; a Longaevus Technologies grant; a Lilly Research Award (28008).

